# Synora: vector-based boundary detection for spatial omics

**DOI:** 10.64898/2026.02.26.708395

**Authors:** Jun-Teng Li, Zhuoran Liang, Zhe Fu, Haojie Chen, Ye-Lin Liang, Na Liu, Qi-Nian Wu, Zekun Liu, Yongqiang Zheng, Jihui Huo, Xiaoxing Li, Zhi-Xiang Zuo, Qi Zhao, Ze-Xian Liu

## Abstract

Tumor–stroma boundaries are critical microenvironmental niches where malignant and non-malignant cells exchange signals that shape invasion, immune modulation and therapeutic response. Spatial omics platforms now resolve these interfaces at single-cell scale, but computational boundary detection remains challenging because heterogeneous neighborhoods can arise either from true compartment interfaces or from unstructured immune infiltration. Here we present Synora, a modality-agnostic computational framework that identifies tumor boundaries using only cell coordinates and binary tumor/non-tumor annotations, making it readily applicable across a broad range of spatial omics modalities. Synora introduces ‘orientedness’, a novel metric that quantifies directional neighborhood asymmetry and distinguishes true boundary cells, where neighbors are spatially segregated by type, from infiltrated regions where cell types intermingle randomly. By integrating orientedness with traditional diversity measures into a unified BoundaryScore, Synora achieves robust boundary identification across synthetic datasets with ground-truth boundaries, maintaining performance under realistic perturbations including 50% missing cells and 25% infiltration. Application to 15 Visium HD spatial transcriptomic datasets across multiple cancer types reveals consistent boundary-enriched gene signatures and cell-type spatial gradients. Validation on a CODEX multiplexed protein dataset demonstrates that Synora’s precise boundary identification enables discovery of clinically relevant cellular neighborhoods and disease-associated spatial patterns missed by frequency-based approaches. Synora enables boundary-aware spatial analyses by making tissue interfaces quantifiable from minimal inputs, helping to standardize interface detection and comparison across spatial omics platforms and biological contexts.

## INTRODUCTION

The spatial organization of cells within tissues is fundamental to understanding complex biological processes, particularly in cancer where the tumor–stroma interface represents a critical zone of cellular interaction and signaling^1,2^. Tumor boundaries are not merely anatomical demarcations but dynamic microenvironments where bidirectional communication between malignant cells and stromal components orchestrates disease progression, immune evasion and therapeutic response^3-10^. Recent advances in spatial omics technologies, including spatial transcriptomics (Visium HD, Xenium, CosMx) and multiplexed protein imaging (CODEX, MIBI-TOF), now enable single-cell resolution mapping of these interfaces^11-16^.

Existing computational approaches have contributed substantially to spatial tissue analysis. Neighborhood-based clustering methods effectively identify regions of cellular heterogeneity and have proven valuable for discovering recurrent microenvironmental patterns across diverse tissue contexts^8,17-19^. Image-based segmentation approaches provide morphologically grounded tissue partitioning and have established important frameworks for quantitative spatial analysis when high-quality histological data are available^6,20^. Copy number variation (CNV)-based methods offer biologically principled compartment definitions by leveraging genomic alterations to distinguish malignant from non-malignant cells, providing robust annotations in specific tumor contexts^21^.

However, these approaches face limitations when applied specifically to boundary detection. Neighborhood-based clustering methods, while effective at identifying cellular heterogeneity, cannot distinguish structured tumor–stroma interfaces from disorganized regions of immune infiltration within tumor nests^17^. In both cases, compositional diversity can be high, but the underlying spatial organization is fundamentally different^8,17^. Image-based segmentation approaches require high-resolution histological data, are computationally intensive, and often fail when applied to architecturally complex tissues with irregular or fragmented boundaries^20^. CNV-based strategies, although biologically grounded, are applicable only when CNVs can be inferred. Many spatial omics assays and experimental designs do not provide recoverable CNV information, including imaging-based proteomics, spatial metabolomics and other non-genome-resolving readouts^21^. Most critically, existing methods lack a principled framework for distinguishing cells that are passively infiltrated within a tissue compartment from those actively engaged at a biologically meaningful interface.

Here we present Synora, a computational framework that leverages vector-based spatial statistics to identify tumor boundaries with high precision and minimal input requirements. The core innovation is the introduction of ‘orientedness’, a novel metric that quantifies directional spatial bias to distinguish true boundary cells from regions of random infiltration. By combining orientedness with traditional diversity measures, Synora achieves robust boundary identification using only spatial coordinates and a coarse tumor/non-tumor annotation per cell.

## MAIN

### Synora framework and orientedness-based boundary detection

Synora follows a three-step workflow (Fig. 1a). First, boundary cells are identified from local neighborhoods using a boundary score that integrates neighborhood heterogeneity with directional spatial organization, producing three compartments (boundary, nest and outside). Second, Synora computes a signed distance-to-boundary for each cell, enabling continuous stratification from stroma through the interface into tumor nests. Third, Synora summarizes boundary architecture with shape metrics that quantify interface abundance and irregularity, providing sample-level descriptors for cross-sample comparisons.

**Fig. 1:**
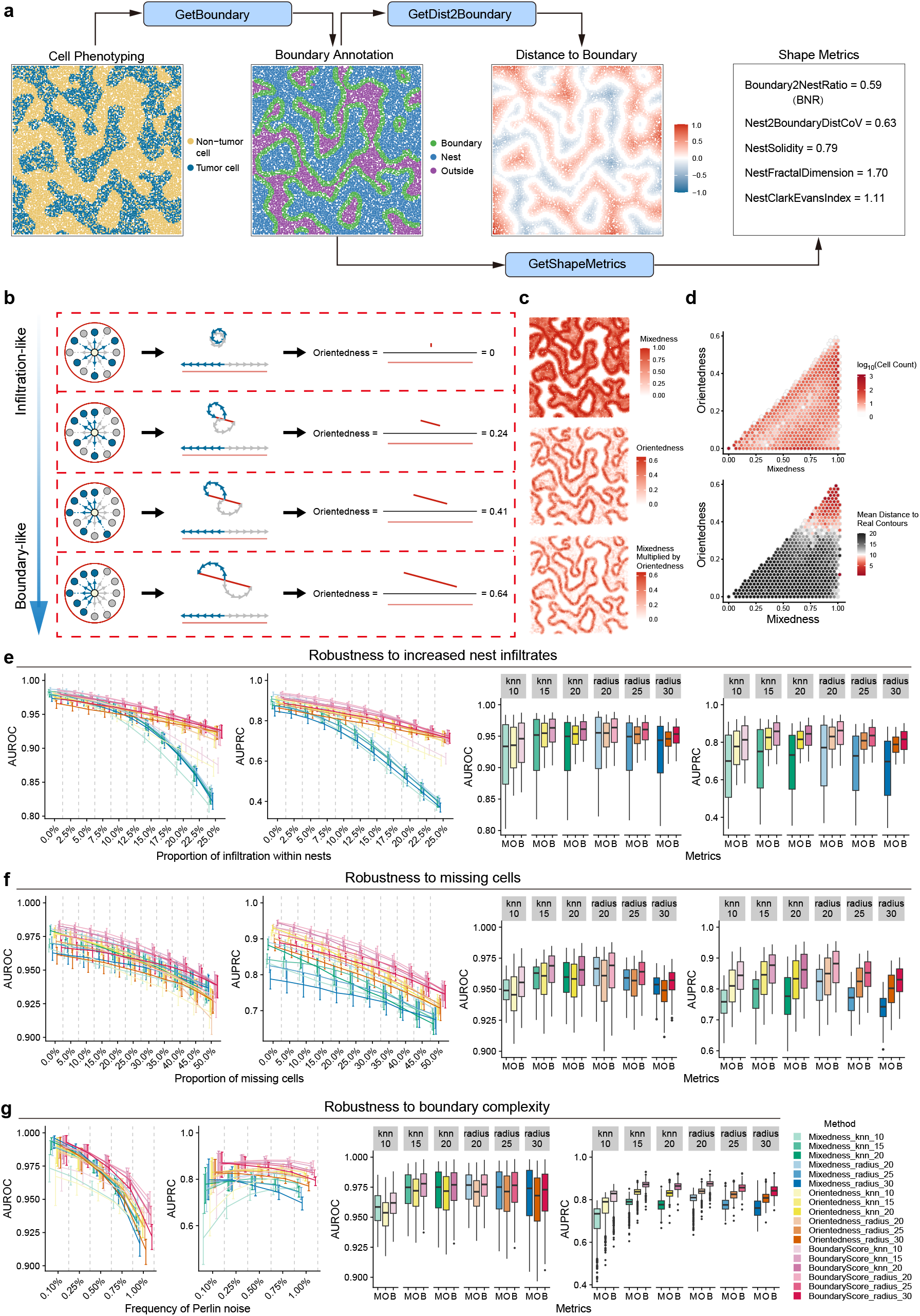
Synora identifies tumor boundaries using vector-based orientedness metric. **a**, Workflow schematic showing Synora’s three main modules applied to spatial omics data. Example visualization shows cell phenotyping (left), boundary annotation (middle) and distance-to-boundary calculation (right) with corresponding shape metrics. **b**, Conceptual illustration of the orientedness metric. Top row shows four scenarios with increasing spatial segregation of cell types (blue = tumor, yellow = non-tumor). Middle row shows vector decomposition for each scenario. Bottom row shows resulting vector sums. Orientedness increases from zero (random distribution) to maximum (complete segregation) whereas Mixedness remains constant. **c**, Application to synthetic Perlin noise dataset. Left, mixedness identifies heterogeneous regions but cannot distinguish boundaries from infiltration. Middle, orientedness captures directional patterns. Right, BoundaryScore (mixedness × orientedness) precisely localizes to ground-truth boundaries (black contours). **d**, Quantitative validation. Left, hexagonal binning shows mean distance to true contours as a function of mixedness and orientedness. High BoundaryScore regions (upper right) exhibit minimal distance to contours. Right, precision-recall curves comparing BoundaryScore to individual metrics across 500 synthetic datasets. **e–g**, Robustness analysis. **e**, Performance maintained under 0–25% infiltration within tumor nests. **f**, Stable performance with 0–50% missing cells. **g**, Consistent accuracy across boundary complexity levels (Perlin frequency 0.001–0.01). Error bars denote the 2.5th–97.5th percentiles across replicates, and the center line represents the median. M, mixedness; O, orientedness; AUROC, area under the receiver operating characteristic curve; AUPRC, area under the precision-recall curve.

A fundamental challenge in spatial boundary detection is distinguishing true tissue interfaces from regions of cellular infiltration. While both exhibit high cellular heterogeneity, they differ critically in spatial organization: at tumor–stroma boundaries, different cell types are spatially segregated, with tumor cells clustering on one side and stromal cells on the opposite side, whereas infiltrated regions show intermixed cell types without directional bias (Fig. 1b, left).

To capture this distinction, we developed orientedness as a metric that distinguishes structured tumor–stroma interfaces from infiltrated regions that are heterogeneous but not directionally organized (Fig. 1b). Synora decomposes boundary evidence into two complementary components. Mixedness captures whether a cell’s local neighborhood contains a mixture of tumor and non-tumor annotations (high for both true boundaries and infiltration). Orientedness captures whether the mixed neighborhood is spatially segregated into opposing directions (high at true interfaces, low when cell types are randomly intermingled). Practically, orientedness is computed by summarizing the directional balance of tumor versus non-tumor neighbors around each cell, with an explicit correction for geometric edge effects so that cells near the imaging boundary are not spuriously favored. Synora then combines mixedness and orientedness into a single boundary score so that only neighborhoods that are both mixed and directionally structured are prioritized as boundary cells.

Using synthetic examples with controlled spatial arrangements (Fig. 1b), we demonstrated the orthogonality of these metrics: as cell distributions transition from random mixing (infiltration-like) to complete spatial segregation (boundary-like), orientedness systematically increases while mixedness remains constant at maximum values. This behavior confirms that orientedness specifically captures directional organization independent of compositional diversity.

### Validation on synthetic datasets demonstrates Synora’s accuracy and robustness

Having demonstrated the orthogonality of mixedness and orientedness through controlled examples, we next evaluated Synora’s boundary-calling performance on synthetic tissues with known ground-truth interfaces generated across a range of boundary complexities using Perlin noise algorithms (Fig. 1c–g, Supplementary Fig. 1). Across simulations, mixedness highlighted heterogeneous regions but did not specifically localize to the interface, whereas the combined boundary score concentrated on cells nearest to the true contour (Fig. 1c,d).

We assessed stability under systematic perturbations, including increasing levels of infiltration-like label noise (0–25% tumor cells reassigned to non-tumor), random removal of cells (0–50% missing data) and higher boundary complexity (Perlin frequency 0.001–0.01) (Supplementary Fig. 2). Performance remained stable under systematic perturbations. Focusing on results using a k-nearest neighbor approach (k = 20), BoundaryScore maintained robustness against increasing nest infiltration (AUPRC from 0.914 to 0.725), missing cells (0.944 to 0.747) and boundary complexity (0.871 to 0.836). In contrast, mixedness alone showed substantially reduced performance, with AUPRC dropping from 0.877 to 0.399 (infiltration), 0.882 to 0.663 (missing cells) and 0.779 to 0.747 (complexity) (Fig. 1e–g, Supplementary Fig. 2). All performance metrics are presented in Supplementary Table 1.

**Fig. 2:**
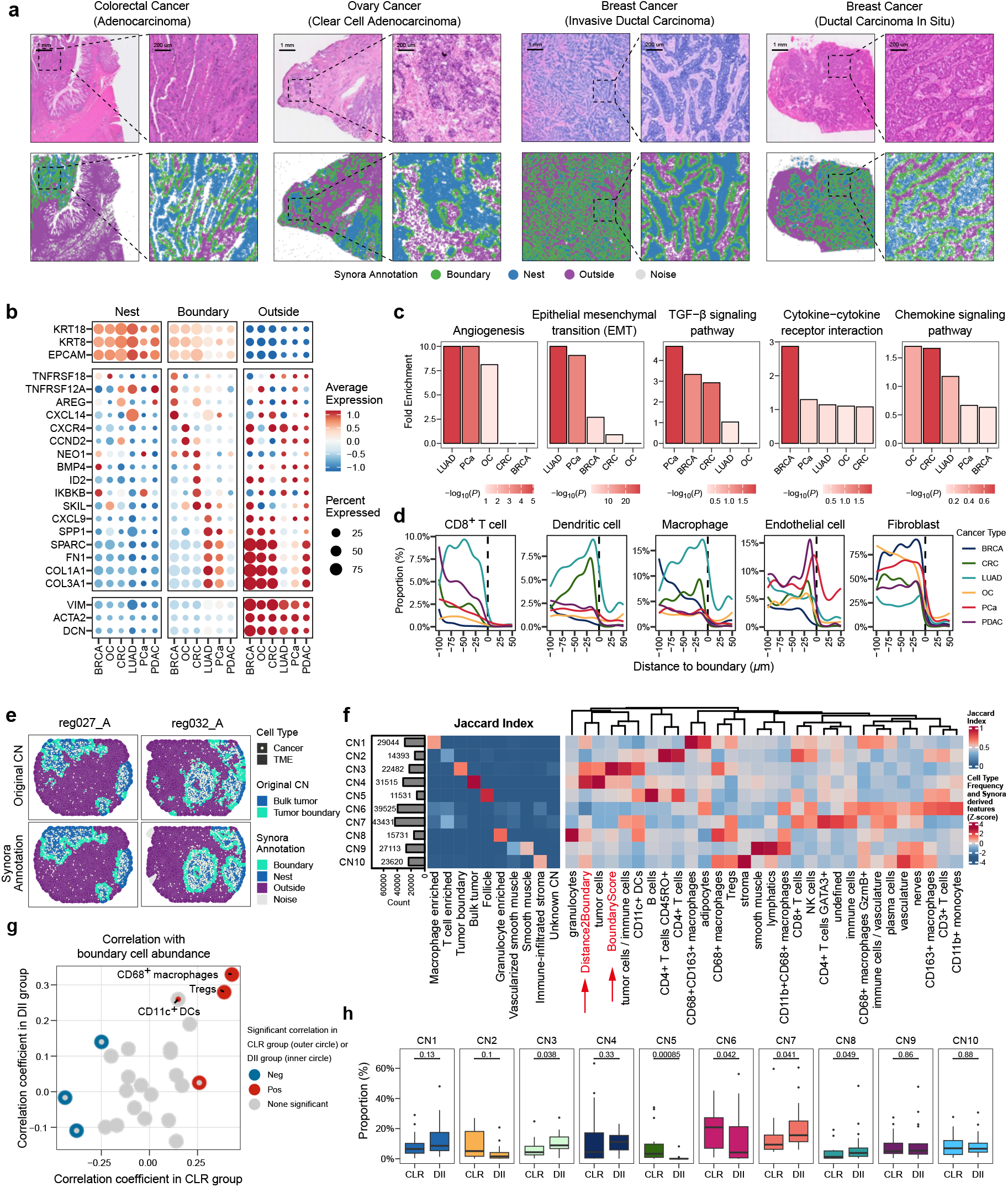
Application of Synora to public spatial transcriptomic and proteomic datasets reveals boundary-enriched molecular signatures and spatial organization patterns. **a**, Representative examples of Synora boundary identification in Visium HD datasets. Top row, H&E histology images. Bottom row, Synora boundary detection. Each image is shown at two magnifications (overview and zoomed region). Scale bars, 200 μm and 1 mm. **b**, Dot plot showing pan-cancer and cancer-type-specific boundary-enriched genes across six cancer types. Dot size represents percentage of datasets showing enrichment; color intensity indicates expression level. **c**, Bar plots summarizing overrepresentation analysis results across cancer types. For each indicated pathway, bars show the fold enrichment of the gene set in each cancer type, whereas the fill color encodes statistical significance. **d**, Distance-binned cell-type composition profiles showing the proportion of different cell types as a function of signed distance to boundary (negative = outside tumor, positive = inside tumor) for each cancer type. **e**, Comparison of boundary identification methods in a CODEX colorectal cancer dataset. Left, original cellular neighborhood (CN) annotation. Right, Synora annotation showing boundary, nest, outside and noise classifications. **f**, Heatmap showing cellular neighborhood (CN) clustering incorporating Synora-derived spatial features (BoundaryScore and distance to boundary). Left panel, Jaccard similarity index comparing original CNs to Synora-enhanced CNs. Right panel, CN composition by cell-type frequency or Synora-derived features (z-scored). Red arrows indicate Synora-derived features. **g**, Scatter plot showing correlation between TME cell types and boundary abundance in patient groups (Crohn’s-like lymphoid reaction (CLR) versus diffuse inflammatory infiltration (DII)). Each point represents a cell type; significant correlations in the CLR group (outer circle) or DII group (inner circle) are highlighted. **h**, Differential abundance analysis of CNs (CN1–CN10) between CLR and DII groups. Box plots show the proportion of each CN in CLR versus DII patient samples. P values from two-sided Wilcoxon rank-sum tests are indicated.

### Synora identifies pan-cancer and cancer-specific boundary-enriched genes in spatial transcriptomic data

To establish biological relevance, we applied Synora to 15 Visium HD spatial transcriptomic datasets spanning six cancer types: breast (BRCA), colorectal (CRC), lung adenocarcinoma (LUAD), ovarian carcinoma (OC), prostate cancer (PCa) and pancreatic ductal adenocarcinoma (PDAC) (Fig. 2a and Supplementary Fig. 3a). Tumor versus non-tumor cells were annotated using established markers, and Synora delineated boundary regions that accurately corresponded to histologically defined tumor–stroma interfaces in matched H&E staining.

Differential expression across nest–boundary–outside compartments revealed a recurrent but cancer type-dependent boundary signature (Fig. 2b and Supplementary Fig. 3b). In subsets of tumors, boundary regions were enriched for extracellular matrix (ECM) and remodeling genes (COL1A1, COL3A1, FN1, SPARC, SPP1), together with more variable enrichment of signaling and activation genes (SKIL, IKBKB, ID2, BMP4, NEO1, CCND2, AREG) and immune and chemokine-associated genes (CXCL9, CXCR4, CXCL14, TNFRSF12A, TNFRSF18).

Consistent with these gene-level patterns, gene set enrichment analysis showed that boundary-associated pathways varied markedly by cancer type (Fig. 2c). Angiogenesis and epithelial–mesenchymal transition were most enriched in PCa and LUAD (with angiogenesis also enriched in OC), whereas TGF-β signaling was highest in PCa (intermediate in BRCA and CRC). By contrast, cytokine–cytokine receptor interaction was most enriched in BRCA, and chemokine signaling peaked in CRC, followed by LUAD. The complete list of boundary-associated pathways is provided in Supplementary Table 2.

To quantify cell-type composition gradients, we stratified cells by distance to the tumor–stroma boundary and calculated cell-type proportions within spatial bins (Fig. 2d and Supplementary Fig. 3c). Across cancer types, tumor cells increased sharply on the tumor side of the interface, whereas fibroblasts were concentrated on the stromal side, reflecting conserved compartmental segregation. Superimposed on this shared architecture, immune and vascular cell distributions showed cancer type-specific boundary enrichment patterns. In CRC, neutrophils were enriched in the stromal compartment, whereas macrophages and dendritic cells were concentrated at the boundary. In PDAC, endothelial cells exhibited a pronounced boundary-associated peak, and in PCa, both Tregs and endothelial cells were elevated near the boundary relative to other tumor types.

### Precise boundary identification with Synora reveals disease-associated cellular neighborhoods in multiplexed protein imaging data

To evaluate Synora’s performance on spatial proteomic data, we applied it to a publicly available CODEX dataset of colorectal cancer^17^ and compared results with the boundary annotations from the original publication. Visual inspection revealed substantial improvements in boundary detection precision (Fig. 2e). As a validation check, distance-binned profiles confirmed that tumor cells were enriched on the tumor side (positive signed distances) and stromal and structural populations on the stromal side (Supplementary Fig. 3d,e).

We repeated the cellular neighborhood analysis from the original study, incorporating two additional Synora-derived spatial features: (i) BoundaryScore and (ii) distance to boundary (Fig. 2f). This analysis identified 10 cellular neighborhoods (CNs), and Jaccard similarity comparisons to the original CN definitions indicated that three CNs were novel (Jaccard index < 0.5). These novel CNs included CN2 (CD4+ memory T cell-enriched), CN6 (T cell–monocyte-enriched) and CN7 (Th2-enriched), representing previously unrecognized microenvironmental niches. Notably, boundary-aware clustering also delineated CN3, a DC–Treg–macrophage-enriched tumor boundary neighborhood.

We next asked whether boundary architecture differed between clinical groups and found that boundary regions were enriched in diffuse inflammatory infiltration (DII) relative to Crohn’s-like lymphoid reaction (CLR) (Supplementary Fig. 3f), a finding not reported in the original study^17^. In DII, and to a lesser extent in CLR, boundary abundance showed positive correlations with macrophages, Tregs and dendritic cells (Fig. 2g).

Differential abundance testing across patient groups revealed five CNs that differed between CLR and DII (Fig. 2h), whereas the original study reported only follicles as differentially abundant between groups. These included CN3 (DC–Treg–macrophage-enriched tumor boundary, P = 0.038), CN5 (follicles, P = 0.00085), CN6 (T cell–monocyte-enriched, P = 0.042), CN7 (Th2-enriched, P = 0.041) and CN8 (granulocyte-enriched, P = 0.049). Together, these findings establish Synora as a robust, platform-independent tool for precise boundary detection that enables discovery of disease-associated cellular neighborhoods.

## DISCUSSION

We developed Synora, a computational framework that resolves a recurring ambiguity in spatial boundary detection by introducing orientedness, a vector-based metric that distinguishes structured tissue interfaces from regions of cellular infiltration. A key strength of Synora is its simplicity: by integrating orientedness with conventional neighborhood heterogeneity measures, it accurately identifies boundaries from minimal inputs (i.e., spatial coordinates and coarse labels), without modality-specific molecular features. This makes Synora modality-agnostic and applicable across transcriptomic, proteomic and metabolomic spatial assays. Furthermore, while we present mixedness and orientedness using binary tumor–non-tumor annotations, the underlying concepts naturally generalize to continuous or probabilistic labels, enabling boundary detection from graded spatial signals such as gene-signature scores, inferred CNV scores, malignancy probabilities or other quantitative molecular phenotypes.

Our validation demonstrates three key advantages over existing methods. First, Synora substantially outperforms scalar heterogeneity metrics by explicitly modeling spatial organization. Frequency-based neighborhood measures cannot separate boundaries from infiltrated regions, and orientedness resolves this ambiguity by quantifying directional spatial bias. Second, Synora is robust to realistic perturbations, including infiltration-like mixing, substantial cell loss and increased boundary complexity. Biologically, these stress tests reflect inflamed tumors with blurred interfaces, clinical samples with imperfect detection or imaging artifacts, and cancers with irregular nest morphology, respectively. Third, unlike image-based segmentation or CNV-based approaches, Synora requires no additional data modalities beyond coordinates and coarse labels, operates efficiently on large datasets and uses a small set of interpretable parameters.

Application of Synora to spatial transcriptomic data across six cancer types revealed both pan-cancer and cancer-specific boundary signatures. The consistent enrichment of ECM remodeling genes (COL1A1, COL3A1, FN1, SPARC) and immune and chemokine genes (CXCL9, CXCR4, CXCL14) at tumor–stroma interfaces aligns with established roles of boundaries as sites of stromal remodeling and immune surveillance^7,22^. Cell-type gradient profiling revealed conserved tumor–fibroblast compartmental segregation overlaid with cancer-specific immune and vascular distributions. These findings suggest that while tumor boundaries share conserved structural features, they harbor disease-specific molecular programs that may influence therapeutic vulnerabilities.

Notably, improved boundary definition can change what downstream spatial analyses are able to detect. In the CODEX cohort, incorporating boundary proximity as an explicit feature in neighborhood discovery enabled boundary-aware clustering and revealed clinical group-associated microenvironmental structure. Synora-enhanced clustering identified CNs whose abundances differed between DII and CLR patients, including CN3, CN7 and CN8 in DII and CN5 and CN6 in CLR. Together, these patterns are consistent with a diffuse inflammatory phenotype characterized by heightened immune activity at the tumor–stroma interface, Th2-associated programs and granulocyte recruitment^23-26^. In contrast, CN5 and CN6 were enriched in CLR patients, aligning with more organized lymphoid structures and the potential immunostimulatory role of monocytes^27-29^. More broadly, these observations illustrate how boundary-aware representations can expose clinically relevant differences that may be underresolved by frequency-based neighborhood definitions.

Despite its demonstrated utility, Synora has limitations that suggest future directions. First, while we validated Synora extensively on synthetic tissues and demonstrated consistent patterns in public datasets, we did not benchmark boundary-calling accuracy against bona fide ground-truth interfaces obtained from orthogonal experimental measurements (e.g., expertly annotated histopathology boundaries, laser-capture-based compartment labeling or matched microdissection profiles). Such evaluations would help quantify absolute accuracy and failure modes in diverse tissue contexts. Second, Synora is currently evaluated on two-dimensional spatial coordinates, and dedicated analyses on true three-dimensional spatial datasets (e.g., cleared-tissue imaging or volumetric spatial transcriptomics) remain to be performed^30-33^. Although the underlying concepts extend naturally to three dimensions, practical issues such as anisotropic sampling, z-dependent resolution and three-dimensional neighborhood definition may affect performance and should be systematically assessed.

Synora is named after the Greek term for ‘boundary’, underscoring our focus on quantifying biologically meaningful interfaces in spatial data. This interface-centric perspective extends beyond cancer to many non-tumor settings where boundaries arise from transitions between compartments rather than from a single cell type. Examples include developmental boundaries, tissue zonation (e.g., periportal–pericentral gradients), inflammatory lesions, wound-healing fronts and host–pathogen interfaces^34-38^. In these contexts, boundary-aware features can help separate structured transitions from dispersed infiltration and provide standardized measures of interface abundance and morphology across conditions and technologies. Together, boundary-centric representations can help connect tissue architecture to molecular states in spatial biology, and as spatial omics technologies continue to mature, approaches that explicitly account for tissue architecture are likely to remain important for translating spatial measurements into biological insights and, in some settings, clinical insights.

## METHODS

### Synora algorithm overview

Synora consists of three modular functions: GetBoundary (boundary cell identification), GetDist2Boundary (distance calculation) and GetShapeMetrics (morphological feature extraction). The core algorithm operates on single-cell spatial data with coordinates (*x, y*) and cell-type annotations (binary or continuous).

### Boundary detection algorithm

#### Step 1: Annotation scaling

For binary annotations, tumor cells are assigned value +1 and non-tumor cells −1. For continuous annotations (e.g., gene expression-derived scores), we use a user-specified or estimate a data-driven cutoff to separate low and high values, then linearly rescale the annotation to the range [−1, +1], to preserve sign for directionality.

#### Step 2: Neighborhood definition

For each cell *i*, we identify neighbors using one of three methods: radius-based, k-nearest neighbors (kNN) or a hybrid defined as the intersection of the two. We recommend radius-based neighborhoods for near-uniform spatial sampling and kNN for variable cell density.

#### Step 3: Iterative denoising and nest identification (optional)

To robustly define cell nests and remove small spurious clusters before boundary calculation, an iterative, density-based denoising procedure is employed. This step can be disabled by the user if not required; when bypassed, the mixedness and orientedness metrics are computed directly from the initial scaled annotations (*A*′) rather than denoised, binarized nest classifications.

#### Step 3a: Initial classification

An initial local cell purity score (*p*_*i*_) is calculated for each cell *i*, by averaging the annotation value of its neighbors:

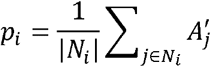

Cells with a *p*_*i*_ score exceeding a user-defined threshold (NEST_SPECIFICITY) are preliminarily classified as ‘Nest’ cells (*N*_*i*_ =1), with all others classified as ‘Outside’ cells (*N*_*i*_ = −1).

#### Step 3b: Denoising iteration

The DBSCAN clustering algorithm^39^ is applied to the spatial coordinates of the ‘Nest’ and ‘Outside’ populations independently. Any resulting cell clusters with a size smaller than a specified minimum (NEST_MIN_SIZE) are considered noise. The classification of these noisy cells is then inverted (*N*_*i*_ → − *N*_*i*_). This denoising process is repeated iteratively until the number of reclassified cells converges to zero, ensuring stable and robust nest definitions. The final denoised classification, *N*_*i*_ replaces the initial scaled annotations 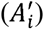 for subsequent boundary calculations.

#### Step 4: Mixedness calculation

For cell *i* with neighbors *N*_*i*_, mixedness is calculated using the scaled annotation formulation. First, compute the mean scaled annotation value of neighbors:

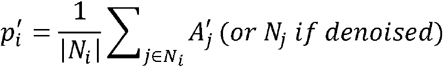

Mixedness is then:

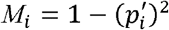

This ranges from 0 (homogeneous neighborhood, where all neighbors have the same annotation) to 1 (maximum diversity, where neighbors are equally split between +1 and −1 annotations). This is mathematically equivalent to the Gini–Simpson index for binary cases^40^. The scaled formulation allows generalization to continuous annotations.

#### Step 5: Orientedness calculation

For each neighbor *j* of cell *i*, we define the unit vector from cell *i* to *j* as 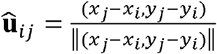. We then calculate the annotation-weighted vector sum: 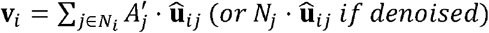. Orientedness is defined as:

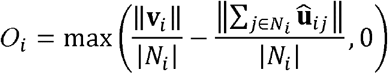

The subtraction term 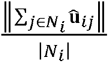, calculated as the unweighted geometric expectation of orientedness, implements an edge correction for cells located at the boundaries of the imaging field or region of interest (ROI). These edge cells have neighbors constrained to one spatial hemisphere, creating artificially elevated orientedness values due to geometric edge effects rather than biological spatial segregation. The max operation ensures non-negative values after edge correction.

#### Step 6: BoundaryScore computation

The final boundary score is: BoundaryScore_*i*_ = *M*_*i*_ × *O*_*i*_ Cells with BoundaryScore_*i*_ above a user-defined threshold are classified as boundary cells.

### Distance to boundary calculation

The GetDist2Boundary function computes a signed distance metric quantifying each cell’s spatial relationship to the boundary interface. The algorithm first partitions cells into ‘boundary’ (cells classified by GetBoundary) and ‘non-boundary’ sets. For each non-boundary cell, we identify its K nearest neighbors (default K = 5) exclusively from the boundary cell population using k-d tree search^41^; using multiple neighbors provides robust distance estimates less sensitive to boundary irregularities. Distance is calculated as the mean Euclidean distance to these K boundary neighbors, with boundary cells assigned distance zero. Finally, distances are signed based on cell annotations: nest cells receive positive values (inside boundary) and stromal cells receive negative values (outside boundary), controlled by the DIRECTION parameter.

### Shape metrics calculation

The GetShapeMetrics function calculates morphological features of tumor nests including: (i) Boundary2NestRatio, perimeter-to-area ratio, quantifying boundary convolution; (ii) Nest2BoundaryDistCoV, coefficient of variation of distances from nest cells to boundary, measuring spatial heterogeneity of nest cell distribution; (iii) NestSolidity, convex hull area divided by actual area, assessing shape compactness; (iv) NestFractalDimension, box-counting dimension^42,43^, quantifying irregularity across spatial scales; and (v) NestClarkEvansIndex, ratio of observed to expected nearest-neighbor distances under random distribution^44^, evaluating spatial clustering versus dispersion of nest cells. These metrics quantify boundary complexity, irregularity and spatial organization.

### Synthetic data generation

We generated synthetic spatial datasets using Perlin noise, a gradient-based noise function that produces fractal-like patterns resembling biological tissue architecture. For each dataset, we constructed a 1,000 × 1,000 coordinate grid and generated a continuous scalar field by applying Perlin noise across multiple frequency settings. We then thresholded the field at zero to define tumor (positive values) and non-tumor (negative values) regions. Cells were placed on a regular lattice with 8-unit spacing and perturbed with Gaussian jitter (σ= 1), and cell annotations were assigned according to the underlying Perlin field value. Ground-truth boundaries were defined as cells lying within 10 units of the zero-level contour, which was extracted using the isoband package. In total, we generated 500 datasets by combining 100 random seeds with five frequency settings.

To evaluate robustness under realistic experimental conditions, we applied three systematic perturbations to the synthetic datasets. First, we simulated immune infiltration by randomly reassigning 0 to 25% of tumor cells to the non-tumor class, mimicking cellular heterogeneity within tumor nests. Second, we simulated missing data by randomly removing 0 to 50% of cells, reflecting segmentation failures or sparse sampling. Third, we varied boundary complexity by adjusting the Perlin noise frequency from 0.001 (smooth, simple boundaries) to 0.01 (irregular, fragmented boundaries), spanning a range of architectural complexity observed in human tissues.

### Spatial transcriptomic data processing and analysis

We analyzed 15 publicly available Visium HD datasets from https://www.10xgenomics.com/datasets (details in Supplementary Table 3). For each dataset, raw sequencing data and H&E images were processed using Space Ranger (version 4.0.1), followed by cell segmentation with the StarDist pipeline. For cell-type identification, single-cell labels were assigned using SingleR. Spatial features and tumor–stroma boundaries were defined using Synora. Cluster-specific marker genes and biological pathways were identified using Seurat and clusterProfiler, respectively. To quantify spatial gradients of cell-type abundance, we employed generalized additive models (GAMs) via the mgcv R package. The occurrence of specific cell types was modeled against the signed distance to the tumor–stroma boundary using a binomial distribution (logit link).

### CODEX data analysis

We obtained processed CODEX data from Mendeley Data (https://data.mendeley.com/datasets/mpjzbtfgfr/1), comprising 140 samples. For each sample, we extracted single-cell coordinates and the provided cell annotations and applied Synora (radius = 70 μm; nest specificity = 0.4; boundary specificity = 0.05) to identify boundary cells and compute BoundaryScore and distance-to-boundary. We performed cellular neighborhood clustering using k-means (k = 10) on local cell-type frequency features augmented with Synora-derived spatial metrics and compared results to the original study’s neighborhoods using Jaccard similarity. Differential neighborhood abundance between disease groups was tested using two-sided Wilcoxon rank-sum tests and group-specific correlations between cell-type abundance and boundary abundance were calculated.

## DATA AVAILABILITY

The Visium HD datasets analyzed in this study are publicly available from 10x Genomics Datasets: https://www.10xgenomics.com/datasets (see Supplementary Table 3 for dataset identifiers). Processed CODEX data were obtained from Mendeley Data: https://data.mendeley.com/datasets/mpjzbtfgfr/1 (140 samples). Derived data supporting the findings of this study are available from the corresponding author upon reasonable request.

## CODE AVAILABILITY

The Synora R package is available at https://github.com/lzxlab/Synora.

## ACKNOWLEDGEMENTS

This work was supported by the Young Talents Program of Sun Yat-sen University Cancer Center (YTP-SYSUCC-0029), the Natural Science Foundation of China (32370698) and the Guangdong Basic and Applied Basic Research Foundation (2023B1515040030).

## Contributions

Z.X.L. conceived the project and provided overall supervision of the study. J.T.L. developed the algorithm and the software, performed analysis on synthetic data, and wrote the manuscript. Z.Liang generated the synthetic data and performed analysis on spatial proteomic data. Z.F. performed analysis on spatial transcriptomic data and prepared the figures. H.C. assisted with figure visualization and reviewed and edited the manuscript. Y.L.L., N.L., Q.N.W., J.H., X.L. and Z.Liu participated in data interpretation and commented on the manuscript. Y.Z., Z.X.Z., Q.Z. and R.H.X. reviewed the manuscript and provided additional supervision. All authors contributed to and approved the paper.

