## Supplementary Figures 1-3 for "Synora: vector-based boundary detection for spatial omics"

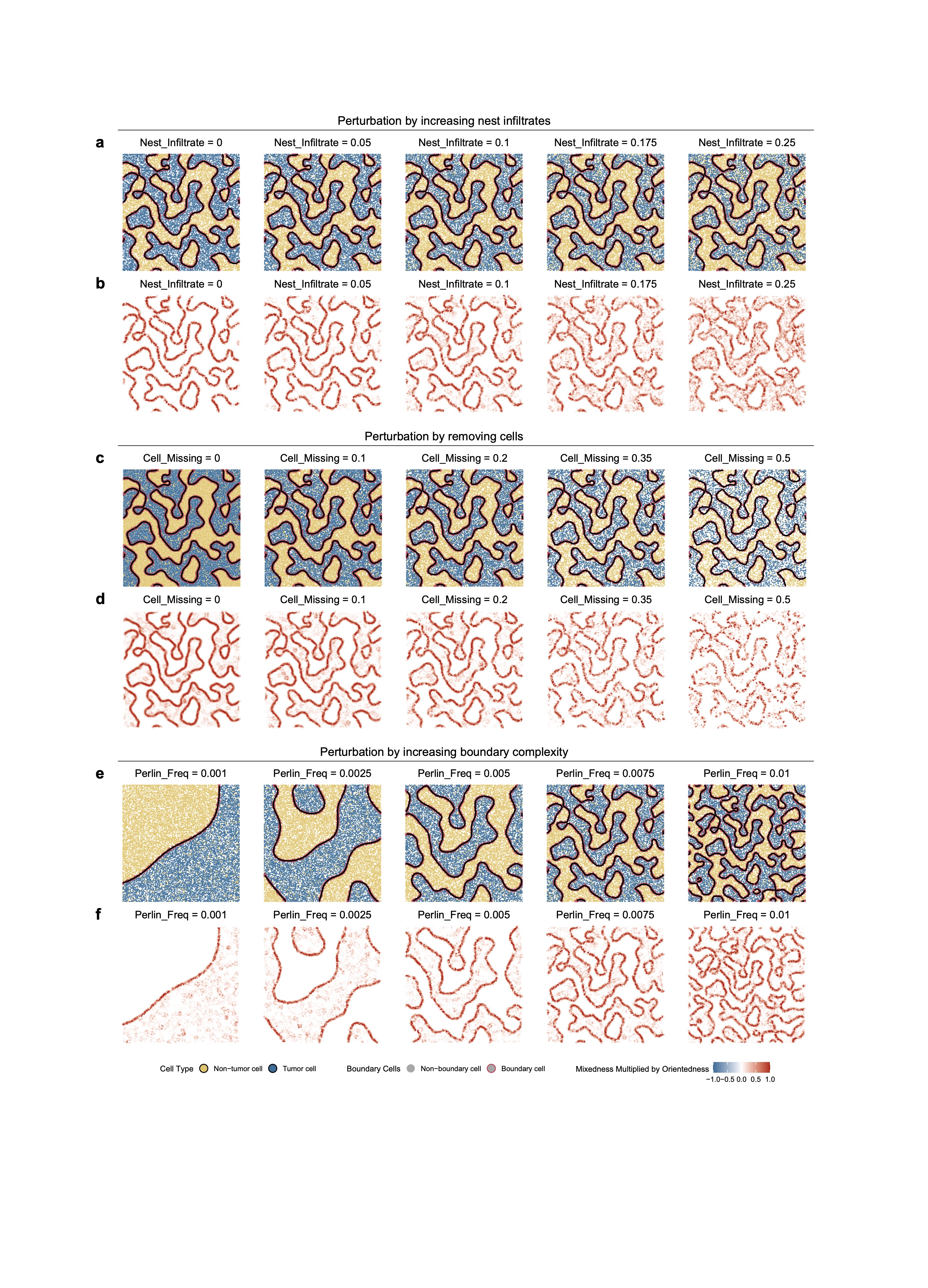

**Supplementary Fig. 1: Visualization of synthetic datasets and Synora scoring under perturbation scenarios.**

**a,b**, Perturbation by increasing nest infiltration. **a**, Spatial distribution of simulated tumor and non-tumor cells with non-tumor infiltration proportions of 0, 0.05, 0.1, 0.175, and 0.25. **b**, Corresponding spatial feature plots of the Synora boundary score (Mixedness × Orientedness). Cells are colored by score values ranging from -1.0 (blue) to 1.0 (red).

**c,d**, Perturbation by removing cells. **c**, Synthetic tissue maps with random cell removal proportions of 0, 0.1, 0.2, 0.35, and 0.5. **d**, Corresponding spatial feature plots of the Synora boundary score.

**e,f**, Perturbation by increasing boundary complexity. **e**, Synthetic tissues generated using Perlin noise with frequencies of 0.001, 0.0025, 0.005, 0.0075, and 0.01. **f**, Corresponding spatial feature plots of the Synora boundary score.

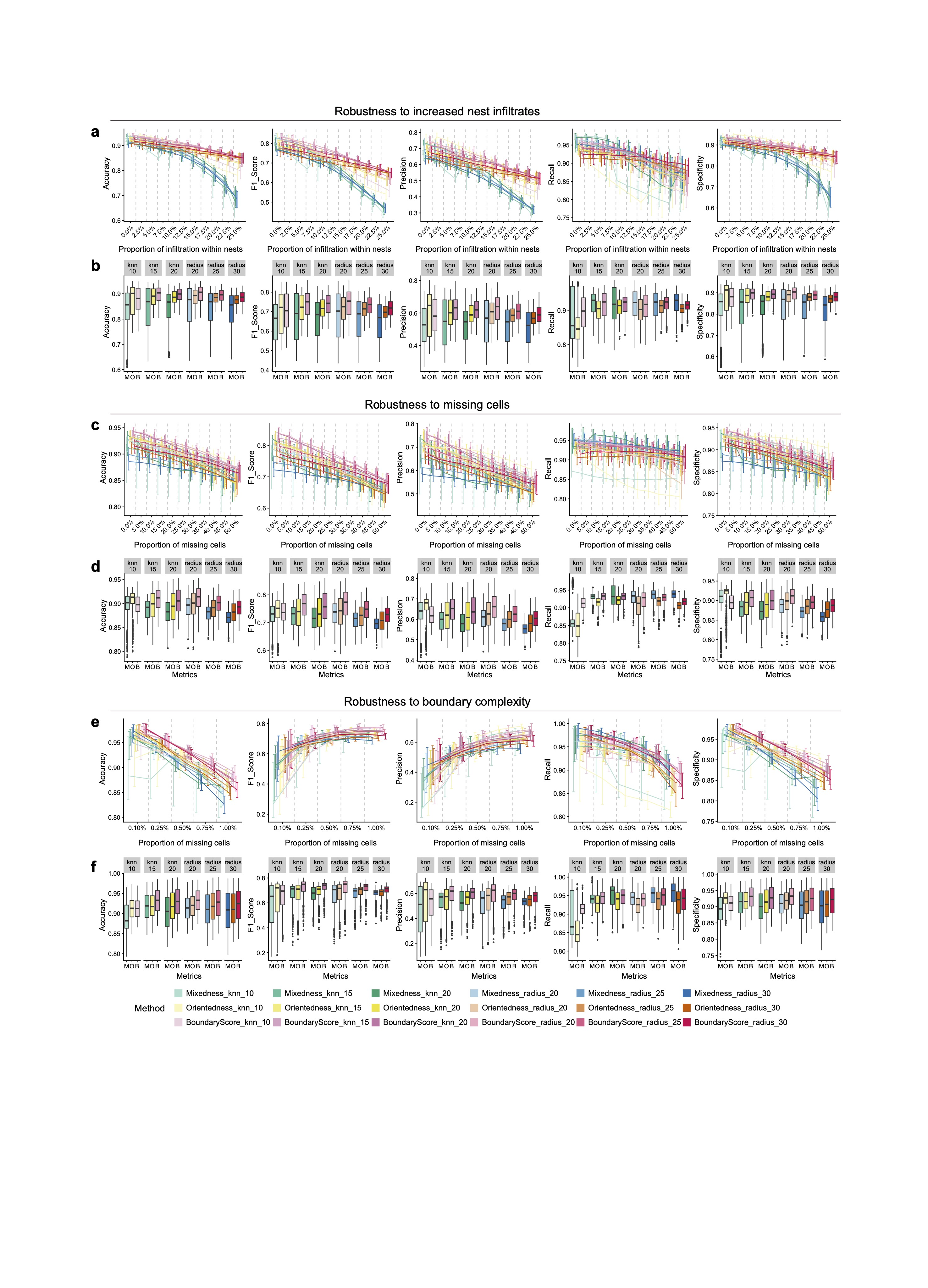
 **Supplementary Fig. 2: Quantitative robustness evaluation of Synora.**

**a,b**, Robustness to nest infiltration. **a**, Five performance metrics (Accuracy, F1, Precision, Recall, Specificity) across six neighborhood definitions (knn: 10–20; radius: 20–30) with increasing infiltration. **b**, Performance metrics comparing Mixedness (M), Orientedness (O), and BoundaryScore (B) components across neighborhood definitions.

**c,d**, Robustness to missing cells. **c**, Performance metrics across neighborhood definitions with increasing cell removal. **d**, Performance metrics comparing M, O and B components under varying cell removal.

**e,f**, Robustness to boundary complexity. **e**, Performance metrics across neighborhood definitions with increasing Perlin noise frequency. **f**, Performance metrics comparing M, O and B components under varying boundary complexity.

Error bars denote the 2.5th–97.5th percentiles across replicates, and the center line represents the median.

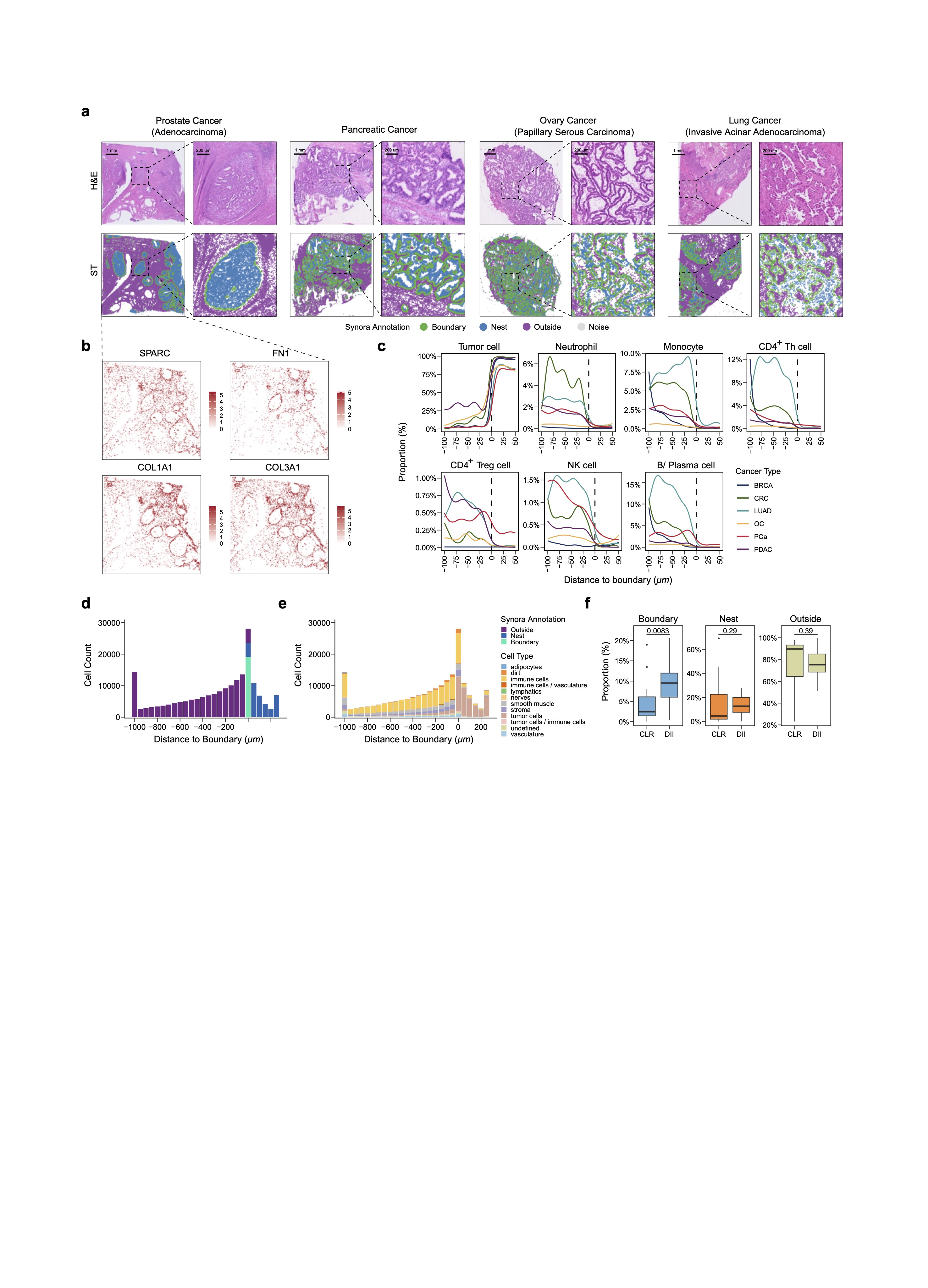
 **Supplementary Fig. 3: Extended application of Synora to Visium HD and CODEX datasets.**

**a**, Application of Synora to Visium HD datasets across four additional cancer types. Top row: H&E histology. Bottom row: Synora spatial domain annotations (Nest, Boundary, Outside, Noise). Cancer types shown are Pancreatic Adenocarcinoma (PDAC), Prostate Cancer (PCa), Ovarian Papillary Serous Carcinoma (OC), and Lung Invasive Acinar Adenocarcinoma (LUAD). Scale bars, 1 mm (overview) and 200 µm (inset).

**b**, Spatial feature plots showing the expression of boundary-enriched extracellular matrix and fibroblast markers (SPARC, FN1, COL1A1, COL3A1) in a representative Prostate Cancer sample.

**c**, Spatial distribution of major cell types relative to the boundary. Line plots display the proportion of specific cell types (Tumor cells, Neutrophils, Monocytes, CD4+ Th cells, CD4+ Treg cells, NK cells, B/Plasma cells) as a function of distance to the Synora-defined boundary (0 µm).

**d–f**, Application to a colorectal cancer CODEX dataset comparing Crohn’s-like lymphoid reaction (CLR) and diffuse inflammatory infiltration (DII) subtypes. **d**, Distribution of total cell counts relative to the Synora boundary in CLR and DII patient groups. **e**, Cellular composition of Synora-defined spatial regions. Stacked bar plots show the proportion of annotated cell types within the Nest, Boundary, and Outside regions. **f**, Comparison of the proportional abundance of the Synora-defined spatial regions between CLR and DII patient samples.
